# Neural correlates of evidence and urgency during human perceptual decision-making in dynamically changing conditions

**DOI:** 10.1101/847756

**Authors:** Y. Yau, M. Dadar, M. Taylor, Y. Zeighami, L.K. Fellows, P. Cisek, A. Dagher

## Abstract

Current models of decision-making assume that the brain gradually accumulates evidence and drifts towards a threshold which, once crossed, results in a choice selection. These models have been especially successful in primate research, however transposing them to human fMRI paradigms has proved challenging. Here, we exploit the face-selective visual system and test whether decoded emotional facial features from multivariate fMRI signals during a dynamic perceptual decision-making task are related to the parameters of computational models of decision-making. We show that trial-by-trial variations in the pattern of neural activity in the fusiform gyrus reflect facial emotional information and modulate drift rates during deliberation. We also observed an inverse-urgency signal based in the caudate nucleus that was independent of sensory information but appeared to slow decisions, particularly when information in the task was ambiguous. Taken together, our results characterize how decision parameters from a computational model (i.e., drift rate and urgency signal) are involved in perceptual decision-making and reflected in the activity of the human brain.

## Introduction

Decisions are often made based on noisy or changing information. A prominent theory in decision-neuroscience, referred to as the evidence-accumulation or drift-diffusion model (Smith and Ratcliff, 2004), posits that deliberation is an integrative mechanism in which information supporting different options accumulates over time until a boundary is reached, at which point the decision is made (Glaze et al., 2015, Gold and Shadlen, 2007, Yang and Shadlen, 2007). Neuroscientific support for the drift-diffusion model comes principally from single-unit recordings in non-human primates. “Accumulator regions” – where neurons exhibit ramp-like increases or drift in their firing towards a decision threshold – have been located in several brain areas within the parietal and prefrontal cortices in a widely-studied dot motion perceptual decision paradigm (Gold and Shadlen, 2007, Hanks et al., 2015, Roitman and Shadlen, 2002, Scott et al., 2017, Shadlen and Newsome, 1996, 2001). This suggests that different pools of selectively tuned, lower-level sensory neurons could feed information to higher-level cortical regions to compute perceptual decisions. However, single-unit recordings provide a spatially narrow view of the brain mechanisms underlying decision-making. Functional magnetic resonance imaging (fMRI) studies have begun to explore the neural substrates of evidence accumulation during perceptual decision-making in humans in attempt to provide a more holistic view (Heekeren et al., 2004, Ploran et al., 2007, Tremel and Wheeler, 2015), but they suffer from poor spatial resolution that makes it difficult to detect brain signals directly related to specific aspects of sensory processing.

Also, some decisions need to be made promptly despite incomplete or changing evidence. Simple drift-diffusion models have difficulty accounting for these situations. A recent theoretical approach suggests that decision-making incorporates an “urgency” signal, independent of the sensory evidence, which grows over time to bring neural activity closer to a decision threshold (Cisek et al., 2009, Mormann et al., 2010, Murphy et al., 2016, Thura et al., 2012). Single-unit recordings in monkeys have implicated the basal ganglia as the neural driver of this postulated urgency signal (Thura and Cisek, 2017). Urgency signals are of interest in human behavior as they may relate to the trait of impulsivity (Carland et al., 2019, Thura and Cisek, 2014, 2016). However, the few studies to date that have employed fMRI to differentiate evidence accumulation and urgency parameters are limited by relatively small sample sizes and univariate BOLD response analysis (Kriegeskorte et al., 2006, Mulder et al., 2014, Park et al., 2014) that may fail to differentiate relevant neuronal populations (Braunlich and Seger, 2016, Gluth et al., 2012).

We designed a novel fMRI task to identify neural substrates of the time-dependent processes that occur during deliberation in a simple sensory decision-making task. We took advantage of the fact that it is possible to reliably decode brain activity related to facial emotion detection. Subjects decided whether a short video of a face presented on screen was transitioning to a happy or sad emotion. Previous work suggests that not only is it possible to decode representation of faces from fMRI signal in extrastriate visual areas (Haxby et al., 2001), but that distinct emotional facial features are uniquely represented in the brain and can be decoded (Kassam et al., 2013, Wager et al., 2015). Our task allowed identification of multivariate patterns indicative of happy or sad faces, which we took to represent the sensory evidence upon which the decision was made. According to evidence accumulator models, information on the upcoming choice decoded from the population of neurons participating in facial processing should increase towards a decision threshold, reflecting the gradual accumulation of evidence in support of the upcoming choice. Analogous to results from single-unit recording studies in nonhuman primates, we hypothesize that decision-making parameters will covary with decoded fMRI activity related to detection of facial emotion. To test the urgency-gating model (Cisek et al., 2009), we included ambiguous trials in the study design. We examined the extent to which neural representation of evidence accumulation contributed to decisions in ambiguous conditions, and whether an urgency parameter improved model fit.

## Results

### Task Design and Aims

This study had three aims. First, we wished to develop a paradigm analogous to the dot-motion task used in primate research that was applicable to human fMRI. This required a visual stimulus for which the information content could be decoded from fMRI. We chose a facial emotion task because others have shown that multivariate pattern analysis (MVPA) could successfully identify the neural correlates of the emotional percept in several brain areas including the fusiform face area. Second, we wanted to measure the flow of information in the brain from perception to action. To do this we trained the multivariate classifier on data from a static task with two facial emotions (i.e., happy, sad). We then applied the same individually tailored classifier to a second task in which subjects were asked to identify emotion in a dynamically changing face (morphing from neutral to either happy or sad). Finally, we wished to test the urgency gating hypothesis; to do this, the dynamic task had two patterns: a gradual unidirectional change and a slower ambiguous pattern. For the ambiguous task, we found that there were two groups of responders: those who responded early, and by definition did so without relevant emotional information, and those who waited for the correct information to accumulate. By comparing these two groups we were able to see if multivariate information drove choice, and whether an urgency-gating signal could explain behavioral performance and brain activity. 53 healthy young participants (age=24.15±5.52, males=24) took part in the present study.

### Multivariate Pattern Analysis of Facial Emotion Detection

A static task was used to localize patterns of brain activity related to each facial emotion using linear support vector machine-learning (SVM) classifiers (Fig 1). Participants viewed almost fully happy or sad faces for 2.5s, then reported the emotional expression via a button box. Beta values derived from a first-level general linear model (GLM) of the BOLD response from each trial (Mumford et al., 2012) were used to search for voxels that carry spatially distributed information about facial emotion – in other words, to find a “happy” or “sad” brain activity pattern. This MVPA approach allows extraction of activity patterns from locally distributed fMRI signal that are more representative of the information processing in an area than simple activation magnitudes. Based on our *a priori* hypothesis and previous studies (Haxby et al., 2000, Wegrzyn et al., 2015), we restricted our analyses to seven regions known to be involved in facial emotion processing resulting in seven classifiers per participant. An association test for the terms “amygdala”, “anterior temporal”, “fusiform gyrus”, “inferior occipital”, “insula”, “intraparietal”, and “superior temporal” was conducted on the Neurosynth meta-analytical database to generate functional masks (Yarkoni et al., 2011). After a k-fold cross validation (*k*=10) to validate the accuracy of the classifiers, each voxel’s distance from the hyperplane (which best splits the two categories of stimuli) was used to determine classifier weight. Subjects who failed quality control or had classifiers that did not significantly decode above chance level were removed from further analysis (n=8). In the remaining group, above-chance classification was possible in all 7 regions of interest (Fig 2).

**Figure 1.**
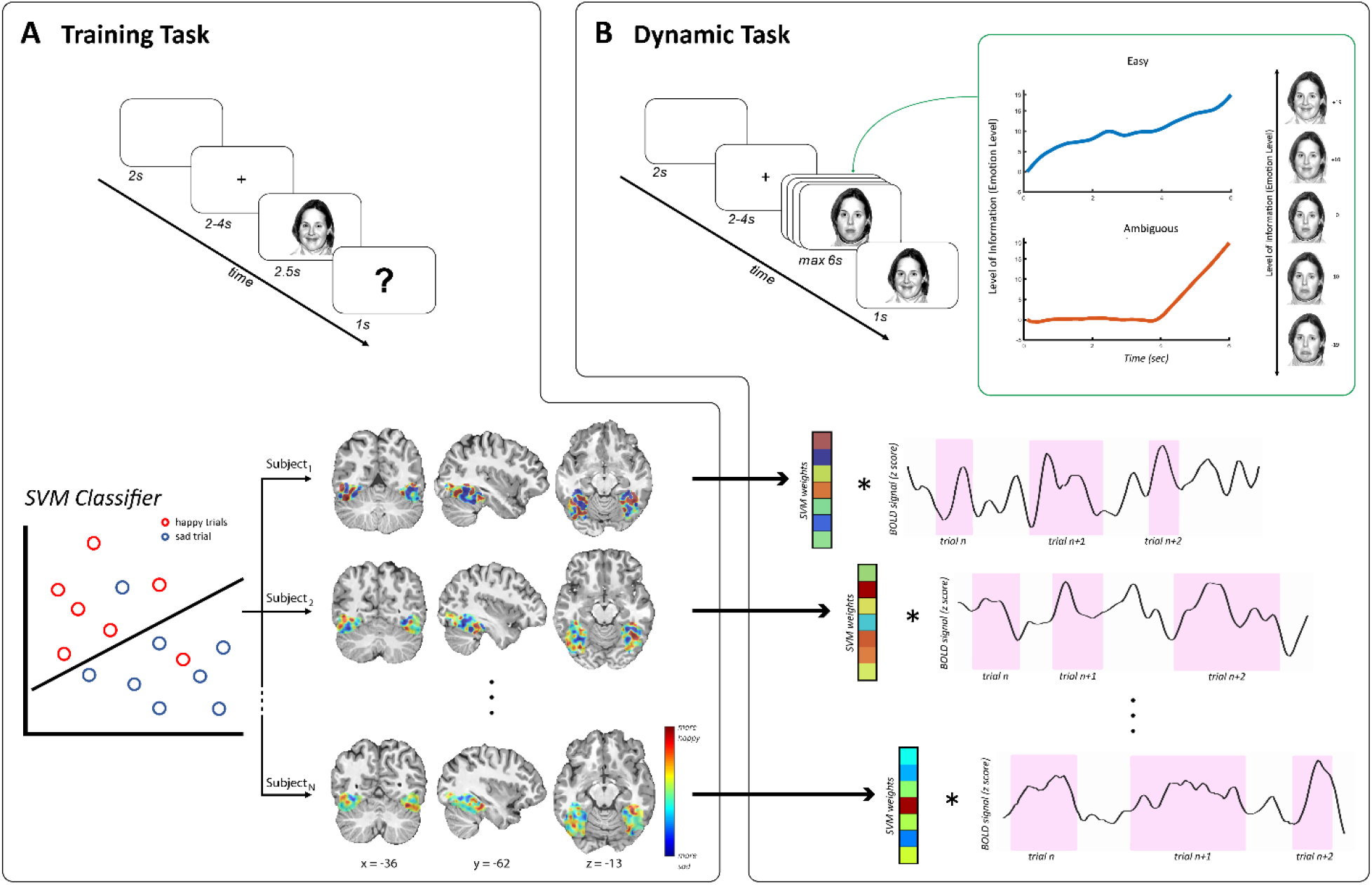
Experiment overview. (A) Training task was used to decode BOLD activity in response to viewing of static happy or sad faces. One classifier was generated for each of 7 regions of interest, per subject. The classifiers were then used to determine support vector machine-learning (SVM) weights, or distance from hyperplane, which were in turn projected to the BOLD activity while viewing faces in the (B) dynamic task. This yielded a neural “code” per trial, per region. Two trial types were used in the dynamic task: (i) easy trials where facial expression gradually morphed towards one of the two emotions and (ii) ambiguous trials where facial expression varied around neutral until two-thirds into the trial after which point emotion rapidly ramped up towards happy or sad.

**Figure 2.**
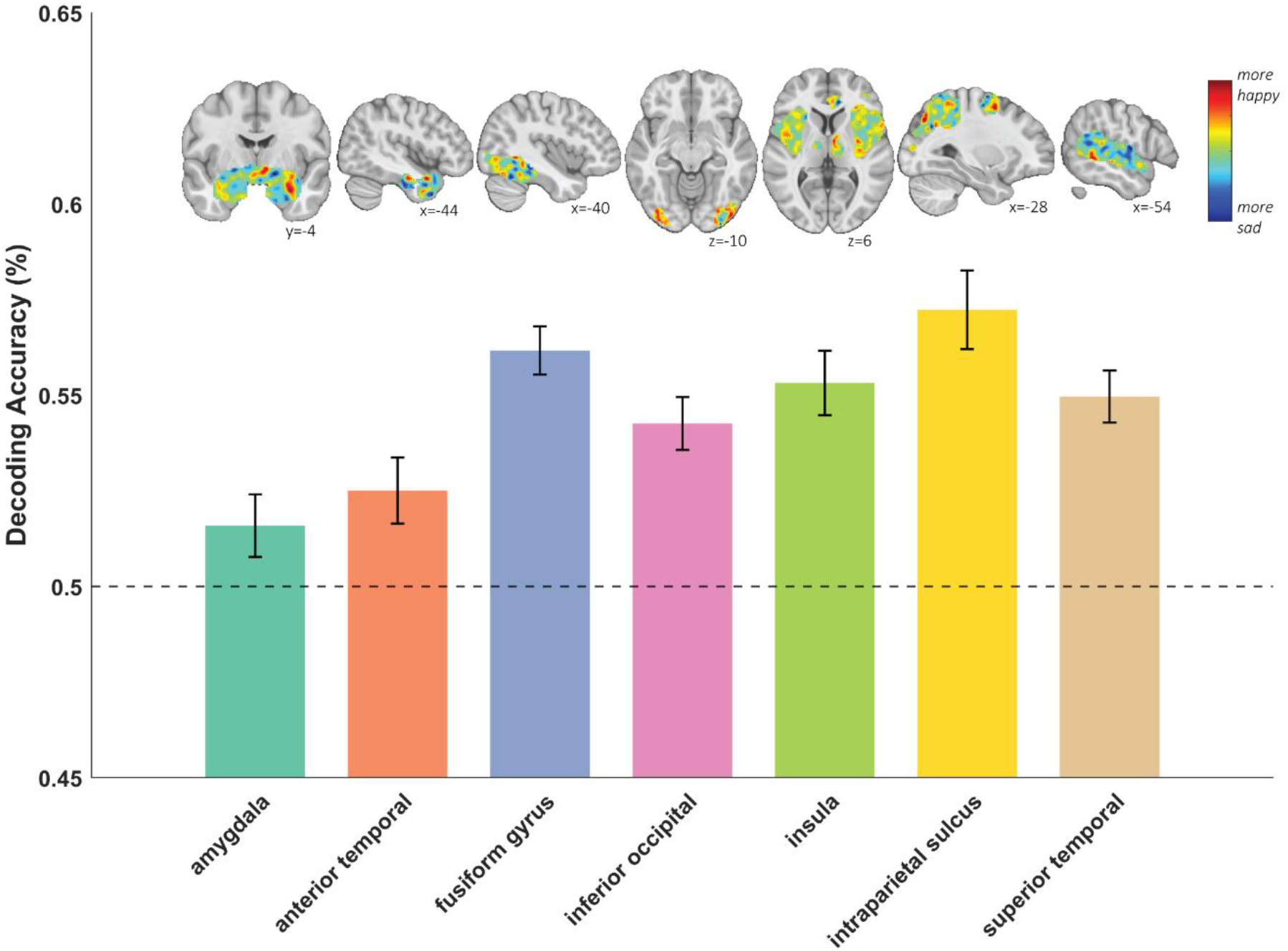
Mean decoding accuracy (n=45) of each region of interest, including amygdala, anterior temporal, fusiform gyrus, inferior occipital, insula, intraparietal sulcus, and superior temporal gyrus. Error bars depict standard error mean. Brain slices show SVC weights from a sample subject with warm and cold colors representing weighting towards happy and sad, respectively.

### Decision Making Task: Behavioral Results

Participants then engaged in a second, dynamic task, which consisted of a video initially showing a neutral face that continuously stepped through intervening morphs to either a happy or sad face over 6s. Participants were instructed to predict, as quickly and accurately as possible, whether the face would be happy or sad at the end of the trial. Within the dynamic task, there were two types of trials modelled after previous work on decision urgency (Thura et al., 2012). Signal (i.e., emotion level) within a given trial was manipulated such that, in one trial type (Easy), the information towards a given emotion gradually increased, becoming easier to discern with time (Fig 1B). In the other trial type (Ambiguous), the information remained ambiguous (close to neutral) throughout the first two-thirds of the trial with information only increasing towards one emotion during the final third of the trial (Fig 1B). Reaction times (RTs) on easy trials (with gradually increasing information) were significantly faster (χ^2^_F_(1,52)=138.77, *p*<.0001) and responses more accurate (χ^2^_F_(1,52)=197.32, *p*<.0001) relative to ambiguous trials (Fig 3). RTs on ambiguous trials were bimodally distributed with responses tending to either be early or late. Overall, subjects also responded faster (χ^2^_F_(1,52)=46.30, *p*<.0001) and more accurately (χ^2^_F_(l,52)=3l.42, *p*<.0001) to trials that were heading towards the happy than the sad direction.

**Figure 3.**
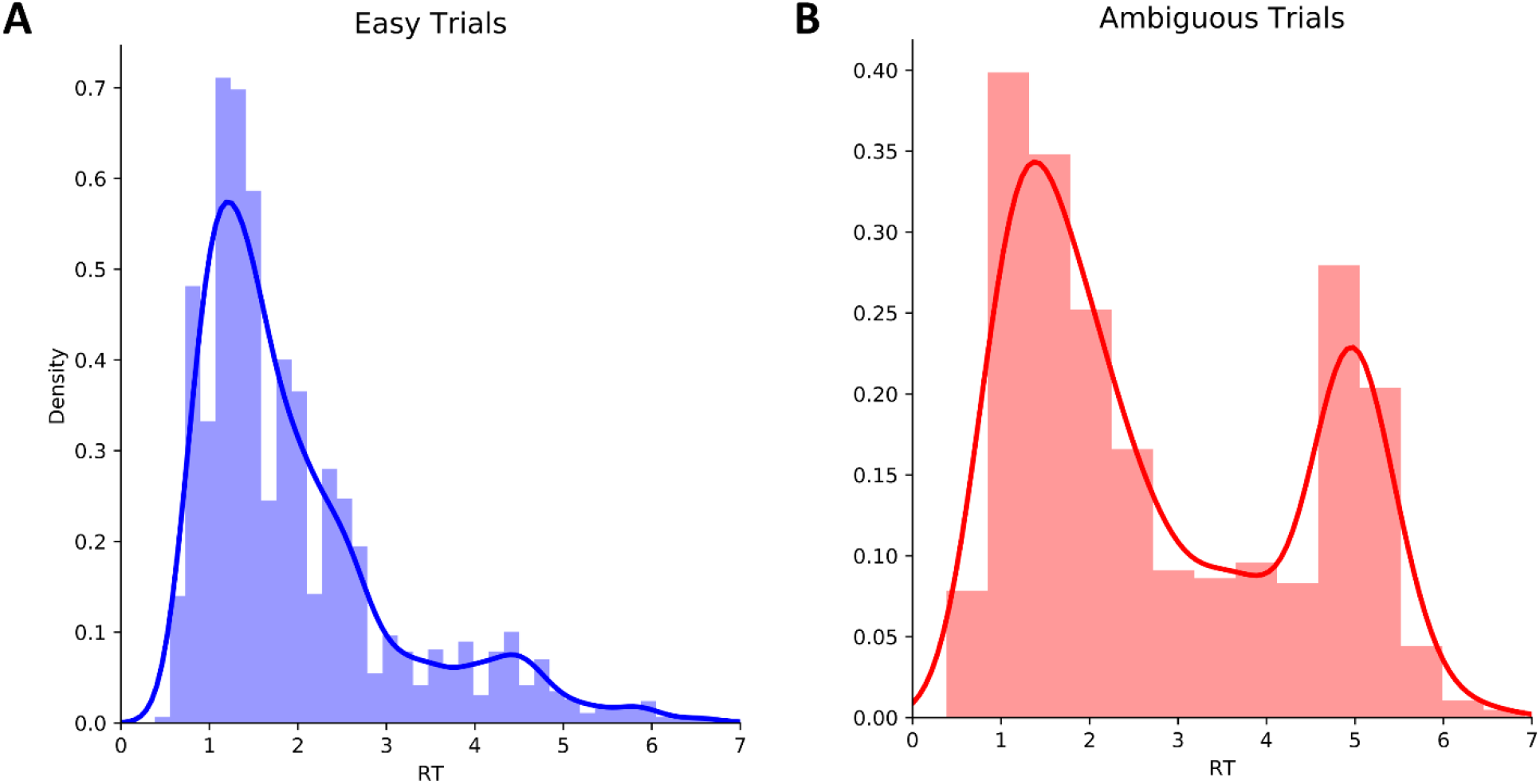
Histogram of reaction time for (A) easy and (B) ambiguous trials. Solid lines reflect the gaussian kernel density estimation.

### Fusiform Code Modulates Drift Rate on a Trial-By-Trial Level

To determine the amount of neural information reflecting perceived facial emotion, SVM weights from the classifiers derived from the static task were projected to the BOLD signal in each trial of the dynamic task to determine the regional MVPA “code” while viewing the morphing video. Combining single-trial regional decoding analysis and drift diffusion modelling (DDM) allowed us to identify context-dependent relationships between single-trial multivoxel BOLD measures and features of decision-making, which are not evident with conventional analyses of reaction times (RTs) and accuracy rates. DDM posits that sensory evidence is accumulated over time until it crosses a decision threshold and the choice is executed. In this framework, behavioral RT distributions are considered to be observations that arise as a function of underlying latent parameters of decision-making. Core parameters include drift rate (*v*), decision threshold (*a*), non-decision time (*t*), and bias (*z*). Here we used a hierarchical extension of the DDM (HDDM) (Wiecki et al., 2013) to estimate decision parameters. This model assumes that parameters for individual participants are constrained by the group distribution but can vary from this distribution to the extent that their data are sufficiently diagnostic. Two central hypotheses were tested using HDDM.

First, we assessed basic assumptions of the model without inclusion of any fMRI data. This involved modulating drift rate by differences in the information available as determined by trial type (analogous to motion coherence in random dot motion tasks (Ratcliff and McKoon, 2008)). High (absolute) drift rates result in faster responses and fewer errors, whereas a drift around zero indicates chance performance with long RT. The drift rate parameter calculated using this basic model was correlated with participants’ overall accuracy in predicting the correct emotion at the end of a trial (*r*=.3331, *p*=.0271), even when RT was used as a covariate in a partial correlation (*r*=.302, *p*=.0463), suggesting that drift rate was a better reflection of behavioral performance than RT alone. Overall, participants were biased towards the happy decision threshold (*z*, mean=0.5606±0.0019).

Second, we tested whether drift rate reflected the regional fMRI MVPA code from our seven regions of interest on a trial-by-trial level (Fig 4A). To test our hypotheses relating multivoxel fMRI activity to model parameters, we estimated posterior distributions not only for basic model parameters, but the degree to which these parameters are altered by variations in neural measures. Regressors were iteratively added to the model to test whether successive additions improved model fit as assessed by the difference in deviance information criterion (DIC) value (Spiegelhalter et al., 2002), with a lower value for a given model (for the whole group) indicating higher likelihood for that model compared to an alternative model, taking into account model complexity (degrees of freedom). Compared to a base model, allowing fusiform MVPA code to modulate drift rate yielded an improved model fit (difference in DIC=26.29) whereas MVPA codes from the other 6 regions did not improve model fit (Fig 4B). Thus, model selection provided strong evidence that trial-by-trial variations in drift rate are modulated by fusiform code as a measure of the evidence for facial emotion. Moreover, while facial emotion is reflected in the entire set of a-priori regions, only information in the fusiform gyrus appeared to influence the decision.

**Figure 4.**
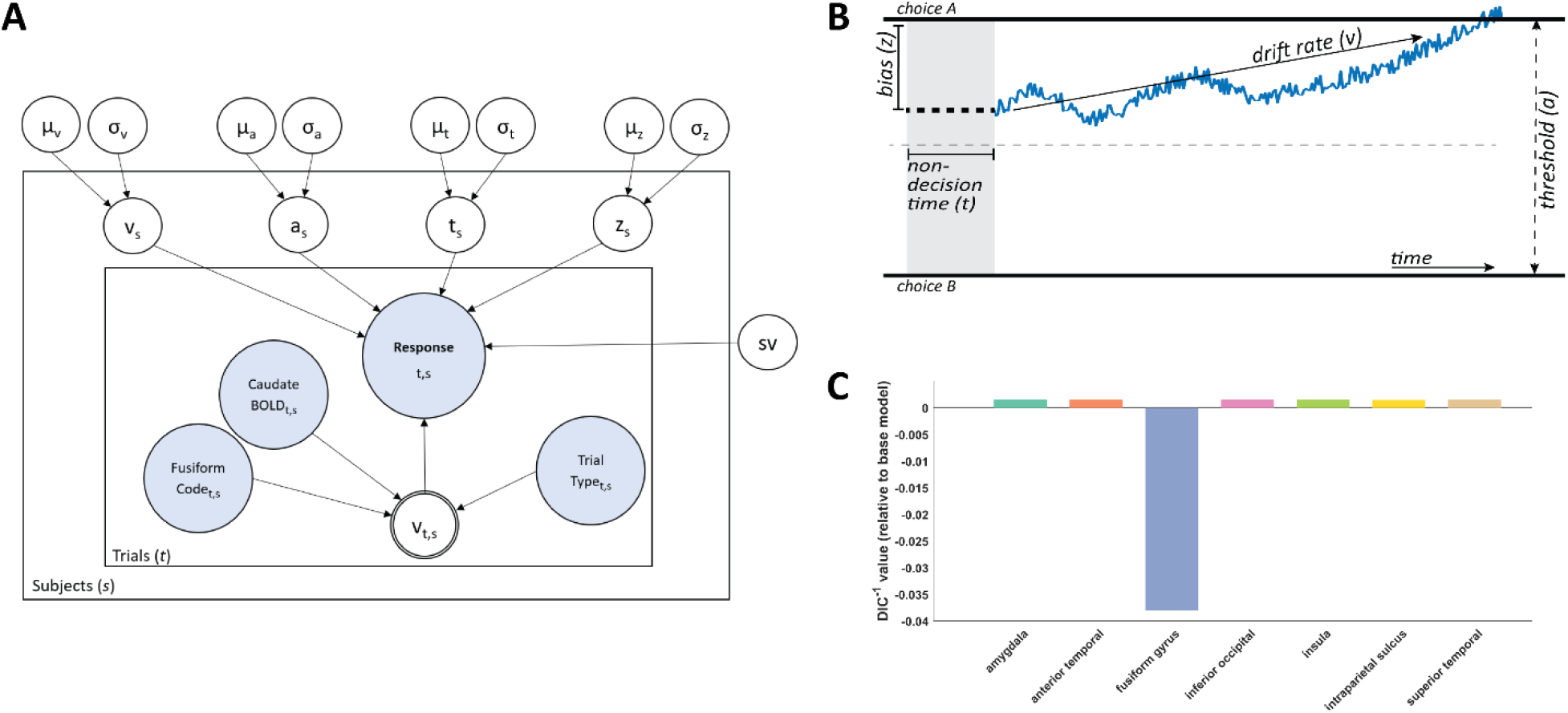
(A) Illustration of Hierarchical Drift Diffusion Model (HDDM) with trial-wise neural regressors. Decision parameters including drift rate (v), decision threshold (a), non-decision time (t), bias (z) and standard deviation of drift rate (sv) were estimated for the group (circles outside the plates with: group mean (μ) and variance (σ)) and subjects (s) (circles in outer plate). Blue nodes represent observed data, including trial-wise behavioral data (accuracy, RT) and neural measures (neural MVPA code from a region as determined by projected SVM weights). Trial-wise variations in v were modulated by neural measures as well as trial type (easy or ambiguous trials). (B) Schematic of the drift diffusion model and estimated decision parameters. Evidence is accumulated over time until one of two decision thresholds is reached at which point a response is made. (C) Model comparison of the seven neural HDDMs. Inverse function of DIC values relative to DIC of the HDDM not containing any neural data are shown (raw DIC values can be found in Supplementary Table 1).

### Neural Circuitry Interacting with Fusiform Code

We were interested in exploring the broader neural circuits that interact with the fusiform face area during perceptual decisions. We used a generalized psychophysiological-interaction (gPPI) analysis (McLaren et al., 2012) to identify brain regions with activity that covaried with the activity of fusiform “seed” voxels as parametrically modulated by the fusiform code. This allows us to identify putative downstream areas that receive the information decoded in the fusiform gyrus in the dynamic task. Two gPPI analyses were conducted with one seed in the left fusiform (center: x=-42, y=-48, z=20) and one in the right fusiform (center: x=44, y=-48, z=-16) (Fig 5, Supplementary Table 2). We found significant increases in connectivity with the bilateral superior and inferior temporal gyrus, lateral occipital cortex, and postcentral gyrus with a right fusiform seed. Similar regions demonstrated increased connectivity with a left fusiform seed but extended to frontal regions including the bilateral inferior frontal gyrus and precentral gyrus as well as the cingulate gyrus and supramarginal gyrus. This asymmetry may be due to use of the right hand for button press.

**Figure 5.**
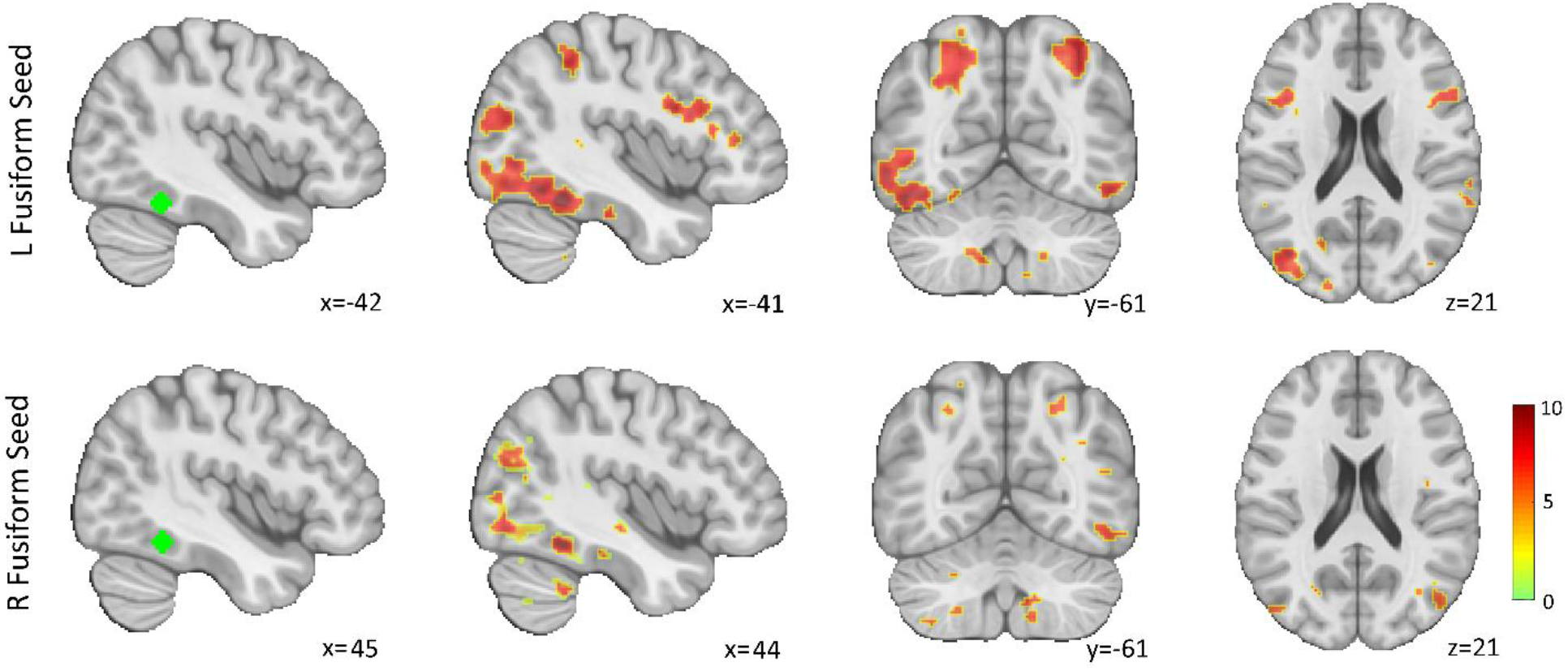
Whole-brain temporal coactivation. Psychophysiological interaction (PPI) from a left (top row) and right (bottom row) fusiform seed as parametrically modulated by the multivariate fusiform code for emotion. Color bar represents t-values.

### Individual Differences in the Tendency to Wait

To further probe the role of fusiform MVPA code, we tested whether the magnitude of this code may differ in easy versus ambiguous trials. In easy trials, the absolute fusiform code significantly differed between correct and incorrect trials (z=2.212, *p*=.0269). This was the not the case in ambiguous trials with no difference in fusiform code observed between correct and incorrect trials (z=-0.103, *p*=.9179). However, a proportion of participants tended to respond rapidly during ambiguous trials, before there was enough information to arrive at a decision. To disentangle these individual differences, we conducted a post-hoc analysis comparing subjects who tended to respond when no information was present in an ambiguous trial versus those who tended to wait for information to be available before responding. Subjects were split into two groups: (1) early responders who, on >=80% of ambiguous trials, responded during the first two-thirds of the trial before information ramped towards one direction (N=21) and (2) the rest, who were categorized as late responders (N=24). As expected, early responders had significantly lower decision thresholds (mean=2.671±0.724) than late responders (mean=5.067 ±0.1.114) in the non-neural HDDM model (*t*(43)=-8.661, *p*<.0001). Early responders were significantly less accurate in predicting trial outcome (mean=53.17%, stdev=0.05) than late responders (mean=77.22%, stdev=13.12) (z=-5.604, *p*<.0001) in ambiguous trials. Early responders had accuracy close to chance in ambiguous trials, suggesting that they were guessing based on partial information rather than making an informed decision.

We next examined the regression coefficients to determine the relationship between trial-by-trial variations in fusiform code and drift rate in a post-hoc analysis (see methods). Our data was split three-ways to generate separate models in HDDM: (1) easy trials across all subjects, (2) ambiguous trials among late responders, and (3) ambiguous trials among early responders. This allowed us to compare drift rates of decisions made during periods of low versus high information. Greater fusiform code increased drift rates in easy trials (95.61% of posterior probability >0) and in ambiguous trials among late responders (97.86% of posterior probability >0). However, this effect was not observed in ambiguous trials among early responders (73.96% of posterior probability >0) (Fig 6). Taken together, our results suggest that fusiform code does not simply drive increases in drift rate, but that this relationship depends on the quality of information as well as individual differences. Early responses during ambiguous trials are made before information is available, therefore the fusiform code cannot affect the response or the modeled drift rate. This further supports the interpretation that the fusiform code is a measure of the evidence that drives the response.

**Figure 6.**
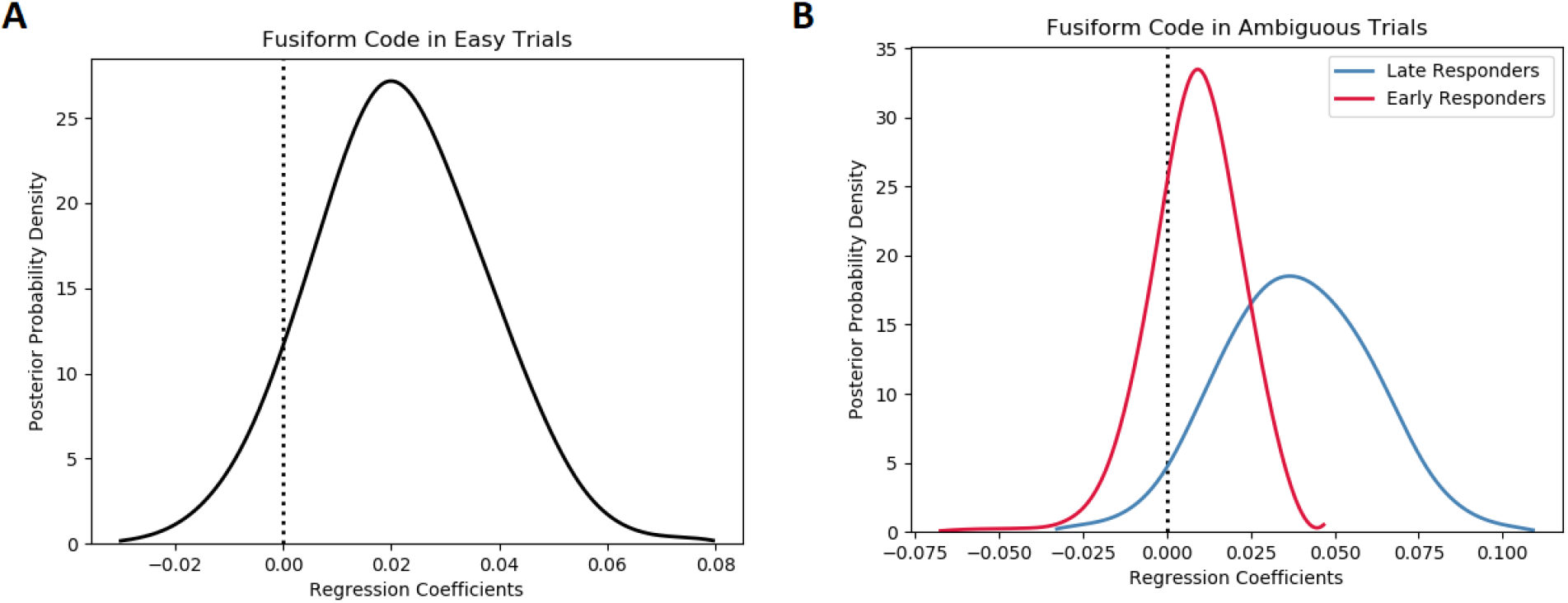
Posterior probability density for modulation of drift rate within (A) easy trials and (B) ambiguous trials split by early (ER) and late (LR) responders. Peaks reflect the best estimates, while width represents uncertainty.

### Caudate BOLD Signal May Reflect Inhibition

We then tried to determine what neural signals differed between late and early responders. A whole-brain group-level GLM revealed that late, versus early, responders had higher caudate activation in two clusters ((1) t=4.45; x=-8, y=4, z=18; (2) t=3.97; x=-8, y=18, z=2) after small volume correction using a structural caudate mask (alpha=.05) as an *a priori* region of interest (Ding and Gold, 2012, Thura and Cisek, 2017) when comparing ambiguous versus easy trails, taking into account the parametric modulation of the fusiform (Fig 7B). This suggests that caudate activity plays a role and may serve to slow down decision in favor of a more accurate choice. Conversely, lower caudate BOLD activity among early responders potentially reflects disinhibition resulting in response prior to having accrued enough evidence.

**Figure 7.**
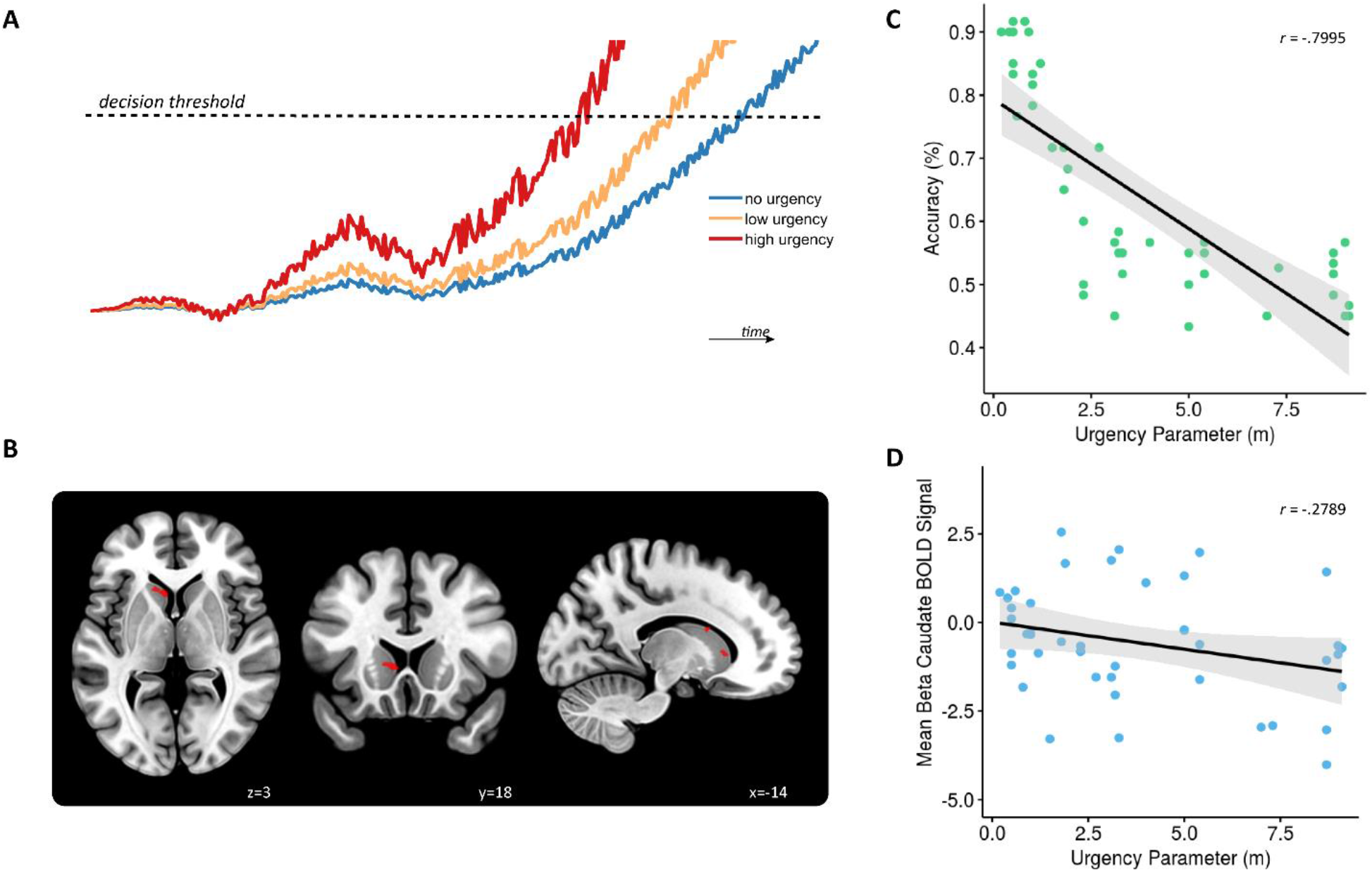
(A) Predicted neural activity y for a sample trial as estimated per the urgency gating model (y(t) = x(t) * u(t)) across time t. This is determined by multiplying a filtered evidence variable x with no (mirroring the drift diffusion model), low, or high urgency u. Both x and u change across time, with u growing as a linear function of time. Once y(t) crosses the decision threshold, a decision is made. (B) A whole-brain group-level general linear model (GLM) revealed that early, versus late, responders had lower caudate activation in two clusters ((1) t=4.45; x=-8, y=4, z=18; (2) t=3.97; x=-8, y=18, z=2), highlighted in red, after small volume correction using a structural caudate mask (alpha=.05) when comparing ambiguous versus easy trails, taking into account the parametric modulation of the fusiform code. An estimated urgency parameter was negatively correlated with (C) performance accuracy among ambiguous trials (r=-.80, p<0001) and (D) mean beta of caudate BOLD signal from within the two significant clusters from our GLM analysis (r=-.28, p=.06).

### Testing the Caudate Signal with the Urgency Gating Model

The caudate is not typically implicated in facial processing (Haxby et al., 2000). Therefore, we sought to test whether its involvement here reflected a not previously described role in facial emotion processing or whether it may be involved in another aspect of decision-making that is independent of the sensory information content, as hypothesized by the urgency gating model. We ran an SVM classifier per participant restricted to the caudate to decode happy and sad faces in the training task. We found that, as opposed to the fusiform and other face processing areas, caudate activity did not decode facial emotions better than chance (mean=50.08%±0.06). Furthermore, we found that adding the trial-by-trial caudate BOLD signal extracted from the aforementioned clusters to the HDDM model did not improve model fit nor did it significantly modulate the drift rate (Supplementary Fig 1). Taken together, this suggests that the caudate did not decode facial information in this task, but rather, perhaps reflects another decision variable untested by the HDDM.

Given the growing literature in support of an “urgency” gating signal (Cisek et al., 2009)(Fig 7A) and the hypothesized role for the basal ganglia in this gating, we tested whether the caudate BOLD may reflect this decision parameter. We used a second model (see methods) that directly tested whether an additional urgency parameter may multiplicatively add to the evidence accumulated, driving it towards a decision threshold, as described by Thura et al. (2012). First, we validated that parameter fits by a non-hierarchical DDM (nDDM) model without urgency corroborated the non-neural basic HDDM results. The estimated decision threshold parameter per participant generated from these two models were highly correlated (*r*=.9461, *p*<.0001). Next, we tested whether a fitted urgency parameter to this nDDM model may relate to the caudate BOLD signal from our clusters. We found that the mean caudate BOLD activity per subject negatively correlated with participants’ urgency parameter with marginal significance (*r*=-. 2789, *p*=.0636) (Fig 7D). Urgency was strongly related to decreased accuracy among ambiguous trials (*r*=-.7995, *p*=<.0001) (Fig 7C). We did not find any significant correlation between urgency and any of our questionnaire measures (i.e., BIS-11, BIS/BAS) (*p*>.05) (Supplementary Table 3). Taken together, our results suggest that caudate BOLD activity reflects a negative urgency signal that dampens participants’ tendency to respond, particularly in ambiguous trials, where there is initially not enough information to make an accurate choice.

## Discussion

Much research on decision making has used simple choice paradigms based on visual evidence, such as dot motion tasks. When used in non-human primates, these tasks allow accurate characterization of the properties of sensory inputs, fitting of computational models to behavior, and identification of neural activity that reflects the underlying sensory evidence or decision variables (Gold and Shadlen, 2007). However, these paradigms are difficult to use in human participants, where trial numbers are usually smaller, and the ability to accurately measure neural activity limited. Here, we took advantage of the large body of knowledge on face processing studied with fMRI, MVPA, and hierarchical Bayesian modelling to overcome these limitations.

We used a dynamic task in which participants had to identify the correct emotion from face pictures that gradually transitioned from neutral to happy or sad. Applying machine learning to fMRI data from a training task, we found patterns of neural activity that encode facial emotion information. We then applied the individual decoders to the dynamic task and showed that the MVPA code in the fusiform gyrus reflected the evidence used to make a choice, as suggested by its relation to computational modelling parameters (drift rate) and by connectivity patterns to areas implicated in sensory decoding, decision-making, and motor control. This suggests that the neural MVPA code was driving decision in our task. Independent of sensory information processing, we observed a stopping (or inverse urgency) signal located in the caudate that appears to delay decisions in ambiguous conditions.

Multivariate encoding of sensory information was found to reflect adjustments of decision parameters in our evidence accumulation model. We confirmed previous MVPA studies by showing that facial emotion can be decoded from each of seven brain regions hypothesized to form the distributed system for facial emotion processing (Haxby et al., 2000, Wegrzyn et al., 2015). However, only the MVPA code from the fusiform gyrus contributed to the drift diffusion model. There was a clear distinction between it and the other 6 regions in terms of DIC (Fig. 4C). This suggests that, while emotional facial features lead to recoverable neuronal activity in the entire face processing network, the fusiform gyrus is central to decoding and feeding the information forward in this decision-making task. These results are keeping with recent evidence that the fusiform gyrus is especially involved in emotion processing (Harry et al., 2013, Wegrzyn et al., 2015). On the other hand, the amygdala, sometimes postulated to specifically decode facial emotion (Haxby et al., 2000), did not influence evidence accumulation in our model. Further support for the role of the fusiform comes from the fact that the strength of the emotional code derived from MVPA was correlated with the estimated drift rate. Single cell recordings in monkeys have shown that drift rate is proportional to the signal-to-noise or coherence of the choice stimulus (Gold and Shadlen, 2007), implying that better sensory evidence is associated with faster accumulation. Note that the relationship between fusiform code and drift rate was contingent on the trial type and on individual differences in participants’ tendency to wait for more information before deliberation. In early responders on ambiguous trials, fusiform code does not contribute to evidence accumulation; this is expected, as there is insufficient evidence in the early portion of ambiguous trials. In sum, our results point to the fusiform gyrus as the key node in decoding facial information for the purpose of this decision-making experiment.

The fusiform gyrus decodes the sensory information, but does it feed this information forward for the purpose of computing a decision variable (Gold and Shadlen, 2007)? We used generalized PPI to identify brain regions where functional connectivity with a seed in the fusiform gyrus was modulated by the fusiform MVPA code. This approach attempts to go beyond simple connectivity to map the actual flow of information used in the task. It has a similar goal to the multivariate pattern covariance methods proposed previously (Anzellotti et al., 2017, Coutanche and Thompson-Schill, 2013) while being rooted in traditional functional connectivity analyses. While our analyses do not reveal directionality, they suggest possible pathways by which information flows to a series of regions belonging to the ventral and dorsal visual streams as well as premotor and cerebellar regions. The connectivity pattern observed here suggests that face information flows to regions implicated in object identification (ventral stream (Mishkin et al., 1983)), action specification (dorsal stream (Goodale and Milner, 1992)), and motor performance. In particular, the information connectivity analysis identified bilateral intraparietal sulcus, an area repeatedly found to encode decision variables in primates (Gold and Shadlen, 2007, Hanks et al., 2006). This information connectivity pattern can be interpreted in the light of the affordance competition model (Cisek, 2007), in which information related to sensory representations and action selection constantly interacts as it moves from occipital to motor areas, and where decisions emerge from a competition between relevant motor outputs. This model predicts that sensory decoding should feed information forward to the medial temporal, parietal and premotor areas involved in converting sensory information into action, as shown here.

The basal ganglia did not display PPI connectivity to fusiform, nor did they appear to encode face information, however they did emerge in our analysis of group differences. Specifically, there was greater caudate activation during ambiguous stimulus viewing in late versus early responders. In the affordance competition model the basal ganglia are thought to bias decisions via cortico-striatal connections (Cisek, 2007, Thura and Cisek, 2017). One type of response bias is to slow down in ambiguous situations, which may reflect negative urgency. Indeed, Cisek et al. (2009) have suggested that the pure evidence accumulation models do not fully account for observed behavior when speed-accuracy trade-offs are present or information is ambiguous. They suggest the presence of an additional model parameter, termed urgency, that is independent of the sensory evidence, but multiplies the drift rate to hasten or slow down decisions when the context demands it. Fortuitously, approximately half our subjects slowed down during ambiguous trials, waiting for the stimuli to morph towards the final emotion, and the others did not. One way for race models to accommodate slower responses is to raise the decision threshold, but because easy and ambiguous trials were intermixed, participants did not have *a priori* knowledge of which type of stimulus would be displayed in any given trial. Another way to account for slower responses on ambiguous trials is lower urgency. Our findings implicate the caudate in slowing down decisions when the evidence is ambiguous. Moreover, the caudate BOLD effect size during ambiguous trials was inversely related to the fitted urgency parameter. (Stated another way, caudate BOLD reflected negative urgency.) These results are consistent with microelectrode recordings in the basal ganglia of monkeys in which the signal was insensitive to evolving sensory evidence but could influence the response speed by modulating activity in sensory processing regions (Thura and Cisek, 2017). The observed caudate activity in our study may reflect the indirect pathway of the basal ganglia – originating from a striatal population of projection neurons thought to generate a net inhibition resulting in a “stopping signal” (Frank and Claus, 2006). For example, in a fMRI study with a dot-motion task, we found that participants slowed their responses when offered the possibility of monetary reward, and that caudate activation during these trials correlated with a raising of the decision threshold (Nagano-Saito et al., 2012). Dopamine signaling was shown to underpin this effect. It should be noted that evidence-independent urgency signals could end up being modeled as drift rate or threshold in evidence accumulation models; to disambiguate urgency from pure evidence accumulation, one needs to dynamically manipulate the amount of information presented, as in the present study (Cisek et al., 2009).

Urgency may underpin the personality trait of impulsivity (Carland et al., 2019). The relation between caudate activation and inverse urgency found here may provide an explanation for a recent meta-analysis reporting hypoactivation of the striatum during reward among individuals with substance use or gambling disorders – two groups often associated with higher impulsivity traits (Luijten et al., 2017). Previous research in the monkey literature also suggests that microsimulation of the caudate nucleus can affect both choice and reaction time (Ding and Gold, 2012). We therefore hypothesized that the urgency signal would relate to impulsive personality, but we did not observe this. Our questionnaire data (i.e., BIS-11, BIS/BAS) may not properly capture motor impulsivity – a personality trait that urgency likely characterizes.

Findings from our study should be considered in light of its limitations. First, both the evidence accumulation and urgency signal are hypothesized to grow in time. Though we used multiband fMRI acquisition to reduce acquisition time, without the ability to record at millisecond resolution, the estimated neural parameters of each model may lack in precision. Second, we used facial emotion as an exemplar of sensory information for perceptual decision-making. Future studies should test whether MVPA decoding can also be applied to other forms of sensory information, and whether the relationship to decision parameters holds. Third, though we observed caudate activity thought to reflect a dopaminergic stopping signal, our study does not measure dopamine nor the indirect pathway *per se.* The implications of this pathway in human decision-making in ambiguous environments merits further research.

In conclusion, by combining model-driven multivariate fMRI analysis, psychophysics, and computational modelling, we characterized two decision parameters underlying human perceptual decision-making processes (drift rate and urgency signal) in the setting of dynamic, changing environments. Our results reveal how these decision parameters are encoded in the human brain and indicate that MVPA techniques can be used to probe and disentangle the biological underpinnings of the decision process. This may be of particular relevance to characterizing brain phenotypes related to disorders of decision-making (e.g., addictions, impulse control disorders, and obsessive-compulsive disorder).

## Acknowledgements

We would like to thank Gaël Varoquaux for their insightful feedback. This research was supported by grants from the Canadian Institutes of Health Research and the Natural Sciences and Engineering Research Council of Canada to A. Dagher. Y. Yau is a Vanier Scholar and received funding from the Canadian Institute of Health Research.

## Author Contributions

YY and AD designed research; YY and MT collected the data; YY, MD, and YZ analyzed the data and contributed methods; and YY, AD, LF, and PC contributed to result interpretation. YY drafted the initial manuscript; all authors contributed to writing of this manuscript.

## STAR Methods

### Contact for Reagent and Resource Sharing

Further information and data requests should be directed to and will be fulfilled by the lead author, Alain Dagher.

### Experimental Model and Subject Details

53 right-handed young, healthy adults (23 males; age 24.02yr±5.49) participated in the present study. Exclusion criteria included current or past diagnosis of a psychiatric disorder, neurological disorder, or concussion, and moderate to severe depression (score >5 on the Beck Depression Inventory (Beck et al., 1961)). All participants gave written informed consent prior to data acquisition and received monetary compensation for their participation. The study was approved by the Montreal Neurological Institute Research Ethics Board.

### Method Details

#### Task Information

Face stimuli were derived from the NimStim database (Tottenham et al., 2009). Photographs of six (3 males) out of 43 models with closed-mouth happy and sad expressions were selected as stimuli for the task because they had the highest identification accuracy in both Tottenham et al. (2009)’s initial validation of the dataset and in our piloting. Face stimuli were made achromatic in MATLAB and presented on a grey background. In order to manipulate the intensity of the emotional expressions, 18 intermediate face stimuli were also generated from the NimStim faces using STOIK MorphMan software (http://www.stoik.com/) to create different emotion levels that gradually transitioned between a model’s neutral and happy or sad face. Thus, emotion levels varied from 0 to 19 in both directions. Two independent tasks were conducted using these stimuli: (1) the training task and (2) the dynamic task.

The static training task (Fig 1A) served to localize patterns of brain activity related to happy and sad faces. Subjects viewed a face with an emotional level >15 for 2.5 secs; this fixed time of display ensured that we eclipsed at least one full TR’s worth BOLD acquisition during fMRI to allow for accurate parameter estimation. After this, a question mark appeared with a maximum time of 1 sec, during which the subject was instructed to respond with their evaluation of whether the face was happy or sad; if no response was made, “Too Slow” was displayed on the screen for 1 sec.

In the dynamic task (Fig 1B), subjects viewed dynamic stimuli of faces “morphing” between expressions. In these trials, a maximum of 60 frames were presented over 6 secs, plus a final image of the correct emotion (with the emotion level > 15) for 1 sec either after a response was made or at the end of a trial if the subject had not yet made a response. Participants were instructed to predict whether the face would be happy or sad by the end of the trial and to respond whenever they felt confident enough to do so. Subjects were asked to respond both as quickly and as accurately as possible. Within the dynamic task, there were two types of trials, namely “easy” and “ambiguous”, which were modelled after previous work (Thura et al., 2012). In both trial types, the first image presented was the model’s neutral face.

In easy trials, all faces presented were of the correct emotion (e.g., a trial in which the correct answer is happy, no sad images are ever presented). Each successive frame had a 65% chance of being one level higher than the previous frame in the direction of the correct emotion. By the final frame, all trials had an emotion level >16. The final frame was presented for 1 sec as soon as the subject made a response, or it was presented as a 1 sec long additional frame if they had not yet responded. Subjects could respond during this final frame only if they had not yet done so.

In ambiguous trials, the probability of each frame during the first two-thirds of the trials (i.e., up to the 40^th^ frame) had a 50% chance of being one level higher than the previous in the direction of the correct emotion, such that the images generally hovered around a neutral valence. To prevent, for example, many slightly happy images and a few very sad images being presented, the maximum level presented in the correct and incorrect direction before the 40^th^ frame were kept within two levels of each other. Furthermore, the maximum level reached in either direction before the 40^th^ frame was limited to 7. In the final third of the trial, there was a steep increase of level in favour of the correct emotion, with a 95% chance that a given frame would be exactly one level higher in favour of the correction emotion than the previous frame. All trials had a final emotion level > 16. As with the easy trials, this final frame was presented for a duration of 1 sec as soon as a response was made, or as a 1 sec long 61^st^ frame during which subjects could respond if they had not yet done so.

In both tasks (i.e., static training and dynamic), a pause followed by a time-jittered fixation cross preceded each trial. The trials were evenly split between happy and sad (determined by the emotion at the final frame for the dynamic task), with the order of trials randomized in every block. Participants took part in 4 runs for the localizer task and 3 runs for the dynamic task. Both tasks had a total of 120 trials each, divided equally among the runs.

#### MRI Acquisition

Neuroimaging was carried out with a Siemens Magnetom Prisma 3T MRI scanner equipped with a 64-channel head coil at the Montreal Neurological Institute (MNI). High-resolution MPRAGE T1-weighted structural images were first obtained for anatomical localization (TR=2.3s; TE=2.3ms; FOV=240mm; scan matrix=192×256×256; voxel size=0.9mm isotropic). Functional data was then acquired with an echo-planar T2*weighted sequence for blood oxygenation level-dependent (BOLD) contrast (TR=0.719s; TE=30ms; scan matrix=104×108×72; flip angle=44°; FOV=208mm; voxel size=2mm isotropic, multiband acceleration factor=8). Here, we capitalized on multi-band acquisition to help improve temporal resolution, allowing for the potential of multiple data points per trial to better characterize signal change during the decision process

### Quantification and Statistical Analysis

#### Analysis of Behavioral Data

Statistics for this study were conducted in R (R Core Team, 2015) and MatlabR2018b (MATLAB, 2018). Due to low sample size, which may increase vulnerability to spurious outliers, non-parametric tests were used to assess the following subject-level data. Mean reaction times and accuracy were evaluated by a Friedman’s test to compare the effect of trial type and emotion while accounting for runs in each instance. Spearman correlations were used to test for all correlations between task performance (i.e., accuracy and RT) and other metrics of interest (e.g., HDDM decision threshold, UGM urgency signal, questionnaires). Wilcoxon Signed-Ranks tests were used to compare neural activity between incorrect and correct trials whereas Wilcoxon Rank Sum tests were used compare between groups (i.e., early versus late responders – see below). In both cases, z-values refer to Wilcoxon’s z (approximation).

#### MRI Preprocessing

Preprocessing and beta extraction were performed using SPM12 (http://www.fil.ion.ucl.ac.uk/spm/) and Matlab. Signals with >4% intensity change were despiked and corrected using ArtRepair Toolbox (Mazaika et al., 2007). Images were corrected for motion, realigned, normalized to the MNI ICBM152 template (Fonov et al., 2009), and minimally smoothed (6mm FWHM Gaussian kernel). Spatial filtering techniques (such as Gaussian smoothing) have been shown to increase the signal-to-noise ratio (Brants et al., 2011, Hendriks et al., 2017), as well as classification performance in multivariate pattern analysis (MVPA) (Op de Beeck, 2010). One subject was excluded from further analysis after quality control due to excessive motion.

#### Multivariate Pattern Analysis & Fusiform Code

Preprocessed functional data were used as input for run-wise GLM first-level designs yielding one regressor for the event of interest, a second for all other events, and six motion regressors (Mumford et al., 2012), creating one GLM per event. This approach is thought to lead to more representative trial-by-trial estimates of the true activation magnitude. Only the beta value (i.e., parameter estimates or coefficients representing effect size from linear regression) for the event of interest was used for all further analysis in generating a classifier in the localizer task. For the dynamic task, fMRI signal extracted through the canonical GLM (i.e., one GLM per run with regressors for the duration of face presentation, intertrial interval, button press, six additional motion regressors as nuisance regressors, and a constant) implemented by SPM was used for statistical analysis.

#### Generation of Regions of Interest

An association test (FDR-corrected, <0.01) for the terms “amygdala”, “anterior temporal”, “fusiform gyrus”, “inferior occipital”, “insula”, “intraparietal”, and “superior temporal” on the Neurosynth meta-analytical database was conducted yielding one brain map per term indicating the probability that term being used in a study given the reported activation (i.e., P(Term|Activation)) (Yarkoni et al., 2011). To avoid overlaps between our regional masks, voxels in overlapping regions were assigned to the region with the greatest z-score from the reverse inference map derived the Neurosynth search terms. These spatially unique maps were then binarized. A linear support vector machine-learning (SVM) algorithm (C=1.0, L2 penalty, square hinged loss, tolerance=0.0001, max iterations=1,000) was implemented using the scikit-learn package in Python (Pedregosa et al., 2011) to classify happy and sad stimuli from the preprocessed beta images after data normalization. Features were extracted within each of the regional masks, without additional voxel selection. A feature’s (i.e., BOLD signal) distance away from hyperplane determined the SVM weight. A k-fold cross validation (*k*=10) was conducted to test the accuracy of the classifier and reveal voxels where local patterns of activation reliably discriminated between happy and sad faces. After subtracting the activity in the preceding inter-trial period and normalization, the SVM weights from the classifier derived from the localizer task were then projected to each trial’s BOLD signal in the dynamic task to calculate the fusiform “code” during viewing of the morphing video.

Statistical significance of the decoder’s accuracy was tested using permutation of the original data per subject with randomly shuffled class labels of the training and testing data sets before supplying them to the classifier (Mahmoudi et al., 2012, Pereira and Botvinick, 2011). This procedure was done 1,000 times in order to generate a null distribution and was used to test how likely a certain classifier accuracy was to occur by pure chance. Due to exchangeability issues between run – that is the risk of predicting runs rather than class label – labels were only permuted within, rather than across data splits (i.e., within each subject, within each run). P-values were calculated as the proportion of instances where permutated data had equal or higher accuracy than the original decoder accuracy divided by the number of all permutations. Eight subjects with classifiers that did not perform better than chance in any of the regions investigated were excluded from further analysis.

#### Fitting the Hierarchical Drift Diffusion Model (HDDM)

The drift diffusion model (DDM), an established dynamic model of two-choice decision processes (Ratcliff et al., 2016), was fitted to subjects’ reaction time (RT) distributions. The DDM simulates two-alternative forced choices as a noisy process of evidence accumulation through time. The mode implies a single accumulator integrating the sample evidence according to a stochastic diffusion process until the evidence accumulated reaches one of two decision bounds, here for ‘happy’ or ‘sad’. The model decomposes behavioral data into four parameters mapped on to the latent psychological process: drift rate (*v*) for speed of accumulation, starting point (*z*) for a response bias towards one choice, non-decision time (*t*) for stimulus encoding and response execution latencies, and critical decision threshold (*a*) for deliberation.

Here we used a hierarchical extension of the DDM (HDDM) (Wiecki et al., 2013) to estimate decision parameters. This method assumes that parameters for individual participants are random samples drawn from group-level distributions and uses Bayesian statistical methods to optimize all parameters at both the group and subject level. In other words, fits for individual subjects are constrained by the group distribution, but can vary from this distribution. This Bayesian approach for parameter estimation has distinct advantages in robustly recovering model parameters estimates for both individual and group levels than other methods, particularly when the number of trials is relatively small. Moreover, HDDM has been shown to reliably estimate DDM parameters, including regressing effects of trial-by-trial variations of neural signals on decision parameters (Matzke and Wagenmakers, 2009, Wiecki et al., 2013). Bayesian estimates allow for quantification of parameter estimates and uncertainty in the form of joint posterior distribution, given the observed experimental data (Gelman et al., 2013). To account for outliers in behavior that cannot be captured by HDDM (e.g., slow responses due to inattention or fast erroneous responses due to action slips), we removed 5% of the trials at each tail of the RT distribution. Markov chain Monte Carlo sample methods were used to accurately approximate the posterior distribution of the estimated parameters. 5,000 samples were drawn from the posterior to obtain smooth parameter estimates, the first 100 samples were discarded as burn-in. Convergence of Markov chains were assessed by inspecting traces of model parameters, their autocorrelation, and computing the Gelman-Ruben statistic (Gelman and Rubin, 1992) to ensure that the models had properly converged.

Two models were used: one without inclusion of any fMRI data and a second that allowed for trial-by-trial variations in neural activity to modulate decision parameters. To test our hypotheses relating neural activity to model parameters, we estimated posterior distributions not only for basic model parameters, but the degree to which these parameters are altered by variations in neural measures (i.e., fusiform code and caudate BOLD activity). In these regressions, the coefficient weighs the slope of the parameters (defined by drift rate *v* and threshold *a*) by the value of the neural measure on this trial, with an intercept, for example: *v*(*t*) = β_0_ + β_1_*condition* + β_2_*fusiform code*(*t*) + β_3_*condition*(*t*)\**fusiform code*(*t*). The regression across trials allows us to infer the degree to which threshold changes with neural activity. Changes in drift rate relate to RT speed and accuracy.

Modulators, in this case the fMRI-derived neural parameters, were iteratively added in to our model to test whether successive additions improved model fit. Model fit was assessed by comparing each models’ deviance information criterion (DIC) value (Spiegelhalter et al., 2002), with a lower value for a given model (for the whole group) indicating higher likelihood for that model compared to an alternative model, taking into account model complexity (degrees of freedom). A DIC difference of 10 is considered significant (Zhang and Rowe, 2014). DIC is widely used for comparisons of hierarchical models where other measures (e.g., Bayesian information criterion) are not appropriate (Frank et al., 2015, Ratcliff et al., 2016). Parameters of the best model were analyzed by Bayesian hypothesis testing, which examines the probability mass of the parameter region in question (i.e., percentage of posterior samples greater than zero). Posterior probabilities ≥95% were considered significant. Note, this value is not equivalent to *p*-values estimated by frequentist methods but can be interpreted in a similar manner.

#### Psychophysiological-Interaction (PPI)

Generalized psychophysiological-interaction (gPPI) analysis (McLaren et al., 2012) was used to identify brain regions with activity that covaried with the activity of the fusiform “seed” voxels as parametrically modulated by the fusiform code. A 6mm sphere centered at the peak voxel from the z-score map of the “fusiform gyrus” search term from Neurosynth in each hemisphere were used as seeds. Our GLM included regressors accounting for periods corresponding to trials for each emotion (i.e., happy and sad) each parametrically modulated by the fusiform code, with intertrial interval duration, button press, six additional motion regressors as nuisance regressors, and a constant. gPPI regressors were created by deconvolving the seed to obtain an estimated neural signal during perceptual decisions using SPM’s deconvolution algorithm, calculating the interaction with the task in the neural domain, and then re-sconvolved to create the final regressor. Participant effects were then used in a group-level analysis, treating participants as a random effect, using a one-sample t-test against a contrast value of zero at each voxel.

#### Fitting the Urgency Gating Model (UGM)

A filtered evidence variable *x* was derived using the following differential equation:

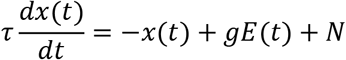

whereby at a given time *t*, the evidence *E* which denotes the amount of information (i.e., emotion level) is multiplied by an attentional fixed gain term *g.* Further, an intra-trial Gaussian noise variable *N* which was defined as six times the signal strength is added. This noise gave us a spread of simulated RT to sufficiently capture the real RT distribution. How far back in time sensory information is considered by the model is determined by the time constant *τ*.

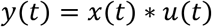

We next computed the estimated neural activity *y.* This was determined by multiplying the filtered evidence with an urgency parameter *u.* A decision is made when the variable *y*(*t*) reaches threshold *T*. A non-decision time of 200ms was added to yield the predicted RT.

Implementation of the non-hierarchical DDM (nDDM) and the urgency-gating model (UGM) differed in two key ways. First, there was no urgency parameter *u* added to the nDDM. In other words, the nDDM assumes that once the variable *x*(*t*) reach the threshold *T*, a decision is made. Second, in the UGM, a low-pass filter of the sensory information in the first-order linear differential equation was applied. The time constant *τ* was set to 200ms for the UGM whereas the maximum trial duration of 6000ms was used as time constant for the DDM. We assumed a time constant of 200ms for the UGM on the basis previous behavioral and physiological studies (Cisek et al., 2009, Thura et al., 2012, Thura and Cisek, 2014). Evidence (*E*), gain (*g*), and noise (*N*) parameters were the exactly same in both models.

In the nDDM, the *T* parameter was adjusted using an exhaustive search to find the variable that minimized the mean squared error between the model’s predicted RT versus the real RT across all trials for each subject. In the UGM, the *u* parameter was similarly searched for using this criterion. Note that for each model, one parameter was adjusted to fit the data; both *T* and *u* influence the means of RT distributions. The models were used to simulate 5,000 trials, the mean of which was used to compare against the real RT distributions.

### Data and Software Availability

Classifier weights from SVM and statistical maps are available for download from Neurovault (see Key Sources Table for further details). Raw data and code used carry out our analysis can be found at https://github.com/yvonnio/face-decoding-fmri. Further details can obtained by request to the lead author.

### Key Resources Table

| REAGENT or RESOURCE | SOURCE | IDENTIFIER |
| --- | --- | --- |
| Deposited Data |  |  |
| Classifier weights (SVM) | This paper | Neurovault: 6025<br>(https://identifiers.org/neurovault.collection:6025) |
| PPI and statistical maps | This paper | Neurovault: 6026<br>(https://identifiers.org/neurovault.collection:6026) |
| Software and Algorithms |  |  |
| MATLAB 2017b | Mathworks | https://www.mathworks.com |
| R | R Core Team | http://www.R-project.org |
| Python 3.5 | Python Software Foundation | http://www.python.org |
| Scikit-Learn | (Pedregosa et al., 2011) | https://scikit-learn.org |
| SPM | Wellcome Department of Cognitive Neurology, London, UK | https://www.fil.ion.ucl.ac.uk/spm |
| HDDM | (Wiecki et al., 2013) | http://ski.clps.brown.edu/hddm_docs |

---

**Supplementary Fig 1.**
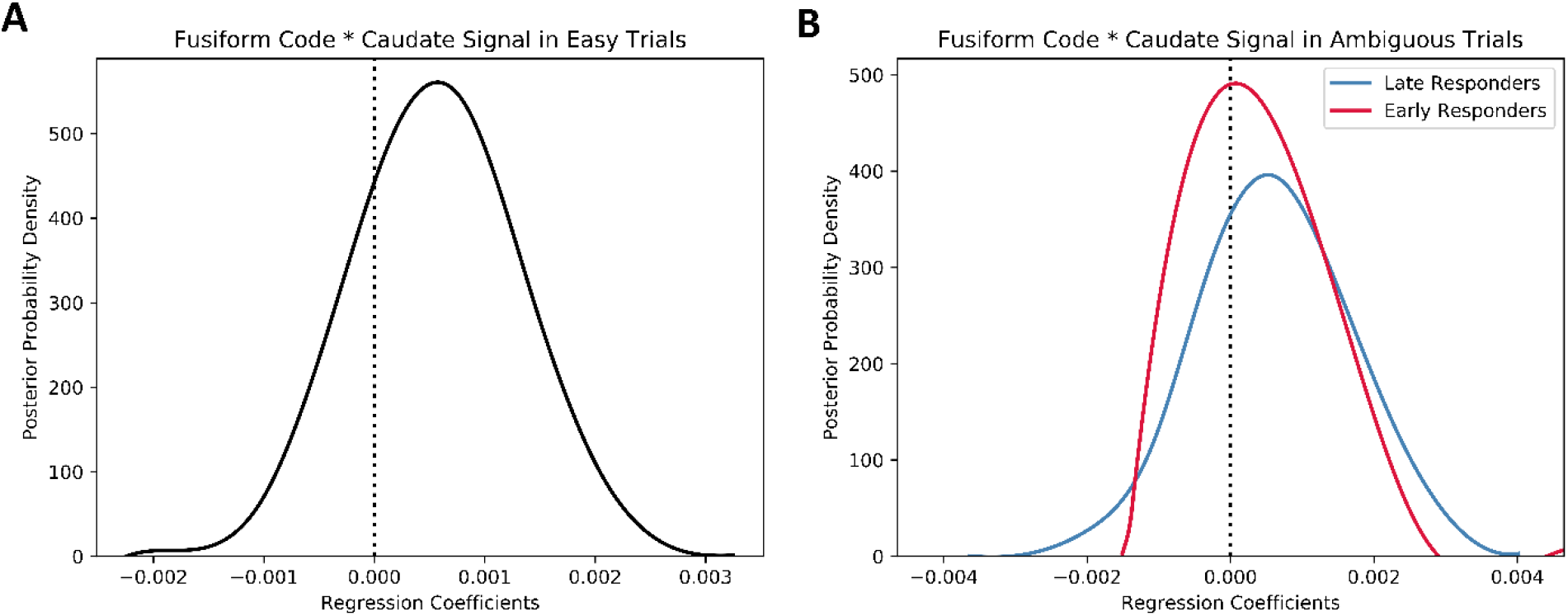
A striatal “urgency” signal may interact with fusiform code depending on task demands. To test this hypothesis, we adapted our HDDM model to assess whether caudate activity, within the significant clusters, could alter decision parameters. As with our HDDM models with fusiform code, we split our data three-ways based on: (1) easy trials across all subjects, (2) ambiguous trials among late responders, and (3) ambiguous trials among early responders. Adding caudate BOLD activity did not improve model fit, as assessed by DIC, beyond a model with only fusiform code. There was weak evidence that the degree to which fusiform code impacted drift rate was modulated by variance in caudate activity in easy trials (79.02% of posterior probability >0) and in ambiguous trials among late responders (71.98% of posterior probability >0). It had little to no effect on ambiguous trials among early responders (54.35% of posterior probability > 0). To test whether caudate may convey information regarding facial emotions, we ran a SVM classifier per participant restricted to the caudate to decode happy and sad faces in the training task. We found that, as opposed to the fusiform and other face processing areas, caudate activity did not accurately decode facial emotions better than chance. Thus, although caudate activity reflected parts of the evidence accumulation process, it did not appear to reflect information processing of facial emotion stimuli nor affect decision parameters as estimated by HDDM.

**Supplementary Table 1.** Raw deviance information criterion (DIC) values comparing HDDM model fit of the seven neural models relative to base.

| Region Name | Difference in DIC value (relative to base) |
| --- | --- |
| Amygdala | 653.49 |
| Anterior Temporal | 650.22 |
| Fusiform Gyrus | -26.29 |
| Inferior Occipital | 662.75 |
| Insula | 672.19 |
| Intraparietal Sulcus | 683.00 |
| Superior Temporal | 668.38 |

**Supplementary Table 2.** Psychophysiological interaction (PPI) from a left and right fusiform seed as parametrically modulated by the multivariate fusiform code for emotion. Related to Figure 5.

| Region | x | y | z | t stat | Number of Voxels |
| --- | --- | --- | --- | --- | --- |
| **L Fusiform Gyrus Seed** |  |  |  |  |  |
| R intraparietal sulcus | 38 | -80 | 32 | 10.79 | 1202 |
| L cerebellum | -14 | -74 | -44 | 10.44 | 267 |
| R cerebellum | 22 | -70 | -44 | 8.26 | 170 |
| R inferior temporal | 54 | -54 | -14 | 10.04 | 551 |
| L fusiform gyrus | -42 | -48 | -20 | 23.96 | 3359 |
| L lateral occipital | -12 | -48 | 54 | 8.03 | 252 |
| R temporal sulcus | 56 | -44 | 4 | 9.45 | 489 |
| L intraparietal sulcus | -48 | -42 | 48 | 9.40 | 360 |
| R intraparietal sulcus | 50 | -36 | 54 | 9.66 | 182 |
| R premotor | 32 | -2 | 52 | 8.99 | 122 |
| L inferior frontal | -42 | 14 | 20 | 9.13 | 335 |
| L inferior frontal | -50 | 34 | 4 | 7.78 | 111 |
| R prefrontal | 48 | 46 | 22 | 11.00 | 143 |
| **R Fusiform Gyrus Seed** |  |  |  |  |  |
| L lateral occipital | -42 | -70 | -2 | 8.86 | 266 |
| R fusiform gyrus | 44 | -48 | -16 | 18.27 | 898 |

**Supplementary Table 3.** Spearman correlations between urgency signal and factors from BIS-11 (Patton et al., 1995) and BIS/BAS (Carver and White, 1994, Heym et al., 2008). No correlations survive significance test (alpha=.05) after a Bonferroni correction for multiple comparison.

|  | Questionnaire | *r* | *p* |
| --- | --- | --- | --- |
| **BIS-11** | Attentional | -0.0751 | 0.6237 |
|  | Motor | -0.2139 | 0.1584 |
|  | Non-planning | 0.0733 | 0.6324 |
| **BIS/BAS** | BAS Drive | -0.1791 | 0.2390 |
|  | BAS Fun Seeking | -0.0456 | 0.7664 |
|  | BAS Reward Responsiveness | -0.3721 | 0.0118 |
|  | BIS Anxiety | -0.0324 | 0.8328 |
|  | BIS FFFS-Fear | 0.0389 | 0.7998 |

## References

1. Anzellotti, S., Caramazza, A., and Saxe, R. (2017). Multivariate pattern dependence. PLOS Computational Biology 13, e1005799.

2. Beck, A., Ward, C., Mendelson, M., Mock, J., and Erbaugh, J. (1961). An Inventory for Measuring Depression. Archives of general psychiatry 4, 561–571.

3. Brants, M., Baeck, A., Wagemans, J., and de Beeck, H.P. (2011). Multiple scales of organization for object selectivity in ventral visual cortex. Neuroimage 56, 1372–1381.

4. Braunlich, K., and Seger, C.A. (2016). Categorical evidence, confidence, and urgency during probabilistic categorization. Neuroimage 125, 941–952.

5. Carland, M.A., Thura, D., and Cisek, P. (2019). The Urge to Decide and Act: Implications for Brain Function and Dysfunction. The Neuroscientist: a review journal bringing neurobiology, neurology and psychiatry 25, 491–511.

6. Cisek, P. (2007). Cortical mechanisms of action selection: the affordance competition hypothesis. Philosophical transactions of the Royal Society of London. Series B, Biological sciences 362, 1585–1599.

7. Cisek, P., Puskas, G.A., and El-Murr, S. (2009). Decisions in changing conditions: the urgency-gating model. The Journal of Neuroscience 29, 11560–11571.

8. Coutanche, M., and Thompson-Schill, S. (2013). Informational connectivity: identifying synchronized discriminability of multi-voxel patterns across the brain. Frontiers in human neuroscience 7.

9. Ding, L., and Gold, J.I. (2012). Separate, causal roles of the caudate in saccadic choice and execution in a perceptual decision task. Neuron 75, 865–874.

10. Fonov, V.S., Evans, A.C., McKinstry, R.C., Almli, C., and Collins, D. (2009). Unbiased nonlinear average age-appropriate brain templates from birth to adulthood. NeuroImage 47, S102.

11. Frank, M.J., and Claus, E.D. (2006). Anatomy of a decision: Striato-orbitofrontal interactions in reinforcement learning, decision making, and reversal. Psychological review 113, 300–326.

12. Frank, M.J., Gagne, C., Nyhus, E., Masters, S., Wiecki, T.V., Cavanagh, J.F., and Badre, D. (2015). fMRI and EEG Predictors of Dynamic Decision Parameters during Human Reinforcement Learning. The Journal of Neuroscience 35, 485–494.

13. Gelman, A., and Rubin, D.B. (1992). Inference from iterative simulation using multiple sequences. Statistical science 7, 457–472.

14. Gelman, A., Stern, H.S., Carlin, J.B., Dunson, D.B., Vehtari, A., and Rubin, D.B. (2013). Bayesian data analysis, (Chapman and Hall/CRC).

15. Glaze, C.M., Kable, J.W., and Gold, J.I. (2015). Normative evidence accumulation in unpredictable environments. eLife 4.

16. Gluth, S., Rieskamp, J., and Buchel, C. (2012). Deciding when to decide: time-variant sequential sampling models explain the emergence of value-based decisions in the human brain. The Journal of neuroscience: the official journal of the Society for Neuroscience 32, 10686–10698.

17. Gold, J.I., and Shadlen, M.N. (2007). The neural basis of decision making. Annu. Rev. Neurosci. 30, 535–574.

18. Goodale, M.A., and Milner, A.D. (1992). Separate visual pathways for perception and action. Trends Neurosci 15, 20–25.

19. Hanks, T.D., Ditterich, J., and Shadlen, M.N. (2006). Microstimulation of macaque area LIP affects decision-making in a motion discrimination task. Nature neuroscience 9, 682.

20. Hanks, T.D., Kopec, C.D., Brunton, B.W., Duan, C.A., Erlich, J.C., and Brody, C.D. (2015). Distinct relationships of parietal and prefrontal cortices to evidence accumulation. Nature 520, 220–223.

21. Harry, B., Williams, M., Davis, C., and Kim, J. (2013). Emotional expressions evoke a differential response in the fusiform face area. Frontiers in human neuroscience 7.

22. Haxby, J.V., Gobbini, M.I., Furey, M.L., Ishai, A., Schouten, J.L., and Pietrini, P. (2001). Distributed and overlapping representations of faces and objects in ventral temporal cortex. Science 293, 2425–2430.

23. Haxby, J.V., Hoffman, E.A., and Gobbini, M.I. (2000). The distributed human neural system for face perception. Trends in Cognitive Sciences 4, 223–233.

24. Heekeren, H.R., Marrett, S., Bandettini, P.A., and Ungerleider, L.G. (2004). A general mechanism for perceptual decision-making in the human brain. Nature 431, 859–862.

25. Hendriks, M.H.A., Daniels, N., Pegado, F., and Op de Beeck, H.P. (2017). The Effect of Spatial Smoothing on Representational Similarity in a Simple Motor Paradigm. Frontiers in Neurology 8, 222.

26. Kassam, K.S., Markey, A.R., Cherkassky, V.L., Loewenstein, G., and Just, M.A. (2013). Identifying Emotions on the Basis of Neural Activation. PLOS ONE 8, e66032.

27. Kriegeskorte, N., Goebel, R., and Bandettini, P. (2006). Information-based functional brain mapping. Proc Natl Acad Sci U S A 103, 3863–3868.

28. Luijten, M., Schellekens, A.F., Kuhn, S., Machielse, M.W., and Sescousse, G. (2017). Disruption of Reward Processing in Addiction: An Image-Based Meta-analysis of Functional Magnetic Resonance Imaging Studies. JAMA Psychiatry 74, 387–398.

29. Mahmoudi, A., Takerkart, S., Regragui, F., Boussaoud, D., and Brovelli, A. (2012). Multivoxel pattern analysis for FMRI data: a review. Computational and mathematical methods in medicine 2012, 961257–961257.

30. MATLAB (2018). MATLAB and Statistics Toolbox Release 2018b. (Natick, Massachusetts, United States: The MathWorks, Inc.).

31. Matzke, D., and Wagenmakers, E.J. (2009). Psychological interpretation of the ex-Gaussian and shifted Wald parameters: a diffusion model analysis. Psychonomic bulletin & review 16, 798-817.

32. Mazaika, P., Whitfield-Gabrieli, S., Reiss, A., and Glover, G. (2007). Artifact repair for fMRI data from high motion clinical subjects. Human Brain Mapping, 2007.

33. McLaren, D.G., Ries, M.L., Xu, G., and Johnson, S.C. (2012). A generalized form of context-dependent psychophysiological interactions (gPPI): a comparison to standard approaches. Neuroimage 61, 1277–1286.

34. Mishkin, M., Ungerleider, L.G., and Macko, K.A. (1983). Object vision and spatial vision: two cortical pathways. Trends in Neurosciences 6, 414–417.

35. Mormann, M.M., Malmaud, J., Huth, A., Koch, C., and Rangel, A. (2010). The drift diffusion model can account for the accuracy and reaction time of value-based choices under high and low time pressure. Judgment and Decision Making 5, 437–449.

36. Mulder, M.J., van Maanen, L., and Forstmann, B.U. (2014). Perceptual decision neurosciences - a model-based review. Neuroscience 277, 872–884.

37. Mumford, J.A., Turner, B.O., Ashby, F.G., and Poldrack, R.A. (2012). Deconvolving BOLD activation in event-related designs for multivoxel pattern classification analyses. Neuroimage 59, 2636–2643.

38. Murphy, P.R., Boonstra, E., and Nieuwenhuis, S. (2016). Global gain modulation generates time-dependent urgency during perceptual choice in humans. Nature communications 7, 13526.

39. Nagano-Saito, A., Cisek, P., Perna, A.S., Shirdel, F.Z., Benkelfat, C., Leyton, M., and Dagher, A. (2012). From anticipation to action, the role of dopamine in perceptual decision making: an fMRI-tyrosine depletion study. Journal of neurophysiology 108, 501–512.

40. Op de Beeck, H.P. (2010). Against hyperacuity in brain reading: spatial smoothing does not hurt multivariate fMRI analyses? Neuroimage 49, 1943–1948.

41. Park, I.M., Meister, M.L., Huk, A.C., and Pillow, J.W. (2014). Encoding and decoding in parietal cortex during sensorimotor decision-making. Nat Neurosci 17, 1395–1403.

42. Pedregosa, F., Varoquaux, G., Gramfort, A., Michel, V., Thirion, B., Grisel, O., Blondel, M., Prettenhofer, P., Weiss, R., and Dubourg, V. (2011). Scikit-learn: Machine learning in Python. Journal of machine learning research 12, 2825–2830.

43. Pereira, F., and Botvinick, M. (2011). Information mapping with pattern classifiers: a comparative study. Neuroimage 56, 476–496.

44. Ploran, E.J., Nelson, S.M., Velanova, K., Donaldson, D.I., Petersen, S.E., and Wheeler, M.E. (2007). Evidence Accumulation and the Moment of Recognition: Dissociating Perceptual Recognition Processes Using fMRI. The Journal of Neuroscience 27, 11912–11924.

45. R Core Team (2015). R: A language and environment for statistical computing. In R Foundation for Statistical Computing. (Vienna, Austria).

46. Ratcliff, R., and McKoon, G. (2008). The diffusion decision model: theory and data for two-choice decision tasks. Neural computation 20, 873–922.

47. Ratcliff, R., Smith, P.L., Brown, S.D., and McKoon, G. (2016). Diffusion Decision Model: Current Issues and History. Trends Cogn Sci 20, 260–281.

48. Roitman, J.D., and Shadlen, M.N. (2002). Response of neurons in the lateral intraparietal area during a combined visual discrimination reaction time task. The Journal of neuroscience: the official journal of the Society for Neuroscience 22, 9475–9489.

49. Scott, B.B., Constantinople, C.M., Akrami, A., Hanks, T.D., Brody, C.D., and Tank, D.W. (2017). Fronto-parietal Cortical Circuits Encode Accumulated Evidence with a Diversity of Timescales. Neuron 95, 385–398.e385.

50. Shadlen, M.N., and Newsome, W.T. (1996). Motion perception: seeing and deciding. Proc Natl Acad Sci U S A 93, 628–633.

51. Shadlen, M.N., and Newsome, W.T. (2001). Neural basis of a perceptual decision in the parietal cortex (area LIP) of the rhesus monkey. Journal of neurophysiology 86, 1916–1936.

52. Smith, P.L., and Ratcliff, R. (2004). Psychology and neurobiology of simple decisions. Trends in Neurosciences 27, 161–168.

53. Spiegelhalter, D.J., Best, N.G., Carlin, B.P., and Van Der Linde, A. (2002). Bayesian measures of model complexity and fit. Journal of the royal statistical society: Series b (statistical methodology) 64, 583–639.

54. Thura, D., Beauregard-Racine, J., Fradet, C.-W., and Cisek, P. (2012). Decision making by urgency gating: theory and experimental support. Journal of neurophysiology 108, 2912–2930.

55. Thura, D., and Cisek, P. (2014). Deliberation and Commitment in the Premotor and Primary Motor Cortex during Dynamic Decision Making. Neuron 81, 1401–1416.

56. Thura, D., and Cisek, P. (2016). Modulation of premotor and primary motor cortical activity during volitional adjustments of speed-accuracy trade-offs. Journal of Neuroscience 36, 938-956.

57. Thura, D., and Cisek, P. (2017). The Basal Ganglia Do Not Select Reach Targets but Control the Urgency of Commitment. Neuron 95, 1160–1170.e1165.

58. Tottenham, N., Tanaka, J.W., Leon, A.C., McCarry, T., Nurse, M., Hare, T.A., Marcus, D.J., Westerlund, A., Casey, B.J., and Nelson, C. (2009). The NimStim set of facial expressions: judgments from untrained research participants. Psychiatry research 168, 242–249.

59. Tremel, J.J., and Wheeler, M.E. (2015). Content-specific evidence accumulation in inferior temporal cortex during perceptual decision-making. Neuroimage 109, 35–49.

60. Wager, T.D., Kang, J., Johnson, T.D., Nichols, T.E., Satpute, A.B., and Barrett, L.F. (2015). A Bayesian Model of Category-Specific Emotional Brain Responses. PLOS Computational Biology 11, e1004066.

61. Wegrzyn, M., Riehle, M., Labudda, K., Woermann, F., Baumgartner, F., Pollmann, S., Bien, C.G., and Kissler, J. (2015). Investigating the brain basis of facial expression perception using multi-voxel pattern analysis. Cortex 69, 131–140.

62. Wiecki, T.V., Sofer, I., and Frank, M.J. (2013). HDDM: Hierarchical Bayesian estimation of the Drift-Diffusion Model in Python. Frontiers in neuroinformatics 7, 14–14.

63. Yang, T., and Shadlen, M.N. (2007). Probabilistic reasoning by neurons. Nature 447, 1075–1080.

64. Yarkoni, T., Poldrack, R.A., Nichols, T.E., Van Essen, D.C., and Wager, T.D. (2011). Large-scale automated synthesis of human functional neuroimaging data. Nature methods 8, 665–670.

65. Zhang, J., and Rowe, J.B. (2014). Dissociable mechanisms of speed-accuracy tradeoff during visual perceptual learning are revealed by a hierarchical drift-diffusion model. Front Neurosci 8, 69.

## Supplementary References

1. Carver, C.S., and White, T.L. (1994). Behavioral inhibition, behavioral activation, and affective responses to impending reward and punishment: The BIS/BAS Scales. Journal of Personality and Social Psychology 67, 319–333.

2. Heym, N., Ferguson, E., and Lawrence, C. (2008). An evaluation of the relationship between Gray’s revised RST and Eysenck’s PEN: Distinguishing BIS and FFFS in Carver and White’s BIS/BAS scales. Personality and Individual Differences 45, 709–715.

3. Patton, J.H., Stanford, M.S., and Barratt, E.S. (1995). Factor structure of the Barratt impulsiveness scale. Journal of Clinical Psychology 51, 768–774.

